## Supplemental Figures for "Activation of a neural stem cell transcriptional program in parenchymal astrocytes"

#### Supplementary figures

Fig. S1

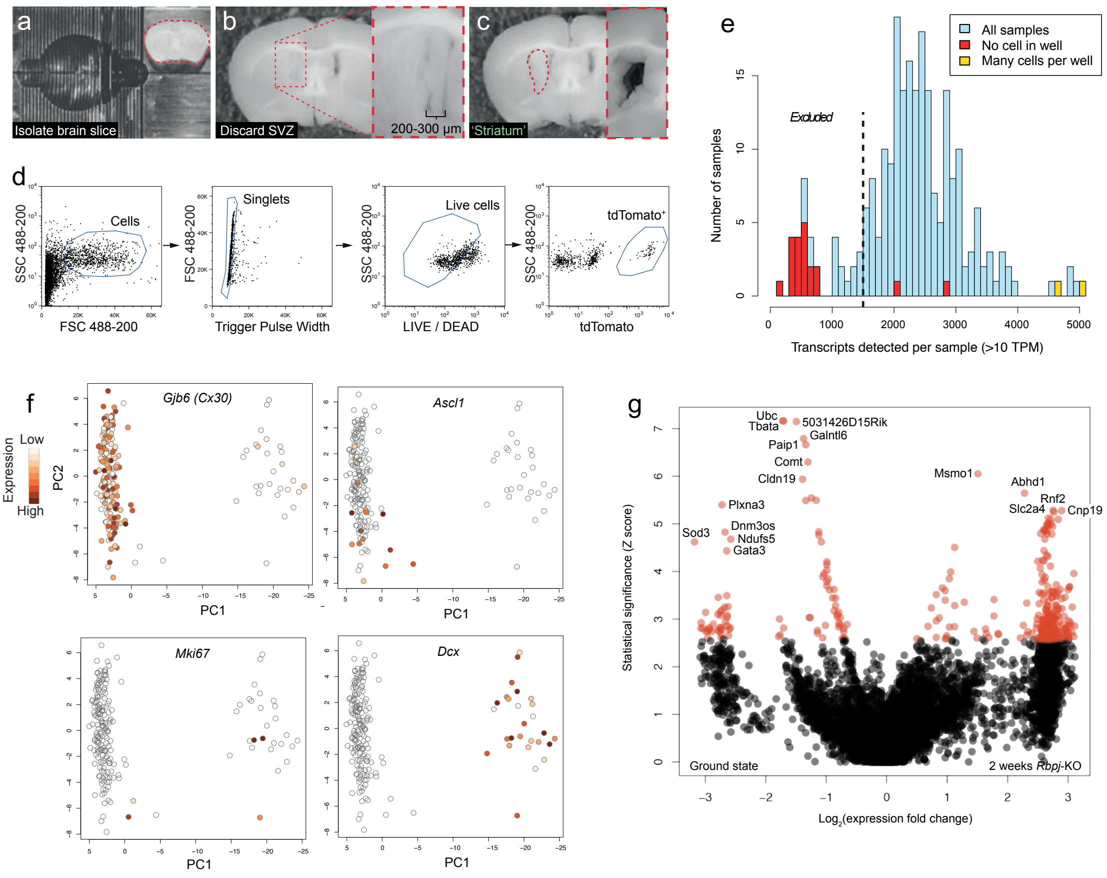


**Figure S1. Isolation of tdTomato$^{+}$ cells for single-cell RNA sequencing and initial analyses.** The dissection procedure for isolation of brain regions is shown in (a-c). First, a ~2 mm thick slice was isolated from the mouse brain (a). The SVZ, including 200-300 µm of the medial striatum, was carefully removed (b) and discarded before the striatum was isolated (c; inset shows region after removal of striatum piece). Striatal cells were sorted based on LIVE/DEAD marker and *tdTomato* expression (d). All wells with no or multiple cells were excluded from further analysis (e). In addition, all samples with <1500 unique transcripts detected were excluded, a conservative exclusion criterion based on the number of transcripts detected in wells into which no cell was sorted (e). Principal component analysis of striatal cells shows the existence of astrocytes (*Gjb6*$^{+}$), activated astrocytes and transit-amplifying cells (*Ascl1*$^{+}$, *Mki67*$^{+}$) and neuroblasts (*Dcx*$^{+}$). Note the relatively small number of transit-amplifying cells (f), probably a result of transit-amplifying cells only existing for a short time, and at only one of our experimental time points. A volcano plot illustrates genes differentially expressed two weeks after *Rbpj* deletion (g). Red data points lie above the statistical significance threshold 2.5.76 (a=0.005).

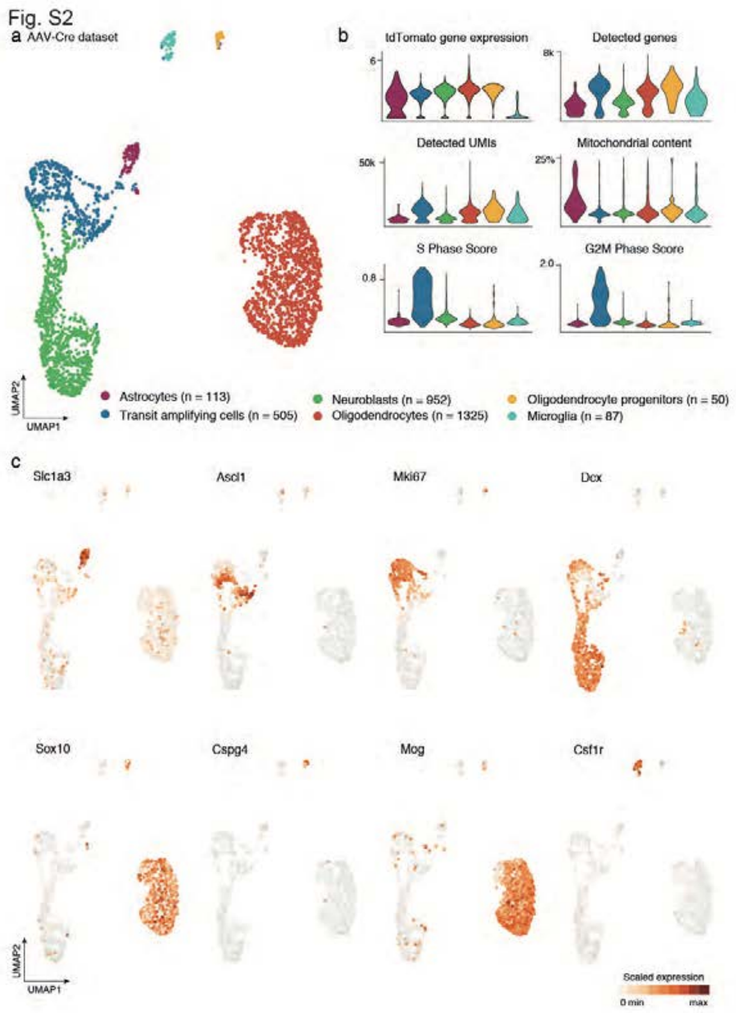


**Figure S2. Cell census from AAV-Cre dataset (10x Genomics single-cell RNA-sequencing experiment) after local recombination of striatal cells.** UMAP visualization indicates the cell types composing the full AAV-Cre dataset (a). This sample encompasses astrocytes and their neurogenic progeny, along with cells from the oligodendroglial lineage, and microglia. Total number of cells detected for each cluster is reported in the legend. Violin plots display the gene expression level of tdTomato, which was used as a means to select cells recombined after viral injection (b). In addition, we report per-cluster numbers of detected genes and UMIs, as well as the percentage of mitochondrial genes, as a measure of quality of the cells. Finally, we report the cell cycle score (S and G2M phase score) to indicate cells, such as transit amplifying cells that are actively dividing. Expression of classical cell type markers are displayed on the UMAP plot (c) to discriminate between populations indicated in (a).

Fig. S3

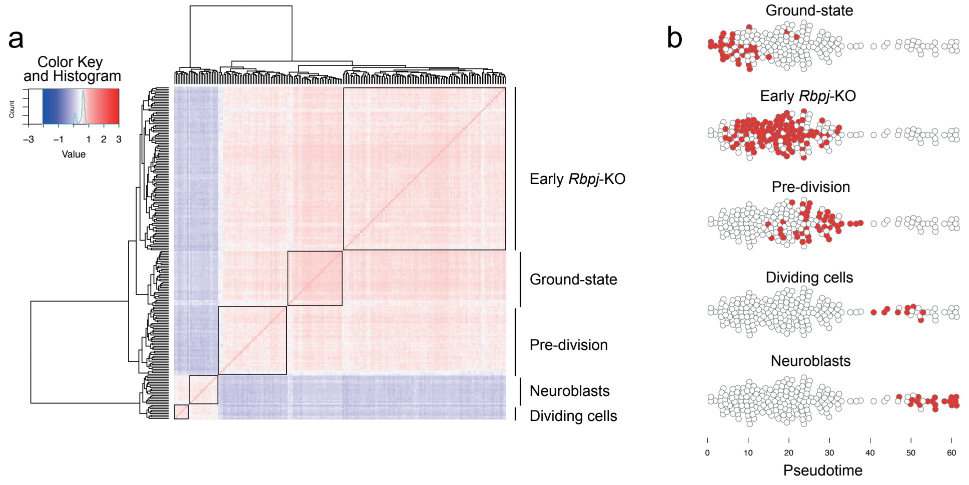


**Figure S3. Hierarchical clustering of striatal astrocytes undergoing neurogenesis**. A correlation heatmap of all 203 striatal cells revealed five cell groups (a). These groups were distributed sequentially along pseudotime (b) and thus likely represent distinct stages along neurogenesis by astrocytes. These groups were used to generate Monocle's gene clustering algorithm (see Fig. 2c): We performed differential expression analysis of each cell group versus the rest, and used the resulting five gene lists as input to the gene clustering algorithm (see Methods).

Fig. S4

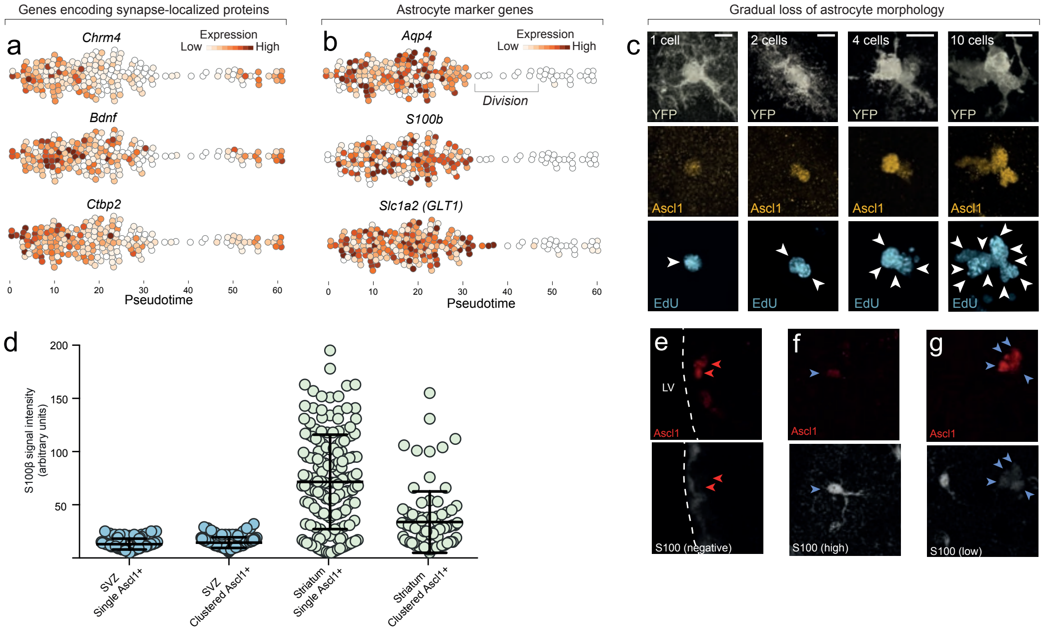


**Figure S4. Astrocytes gradually lose astrocyte features as they enter the neurogenic program.** Genes encoding proteins localized to synapses are downregulated early upon entry into the neurogenic program (a), suggesting that some normal astrocyte functions are immediately compromised as neurogenesis is initiated. Interestingly, these same synapse-associated genes are upregulated again as cells differentiate into neuroblasts, suggesting that they are important for synapse maintenance in both astrocytes and neurons. Common astrocyte marker genes such as *Aqp4*, *S100b* and *Slc1a2*, are maintained in astrocytes approximately until they enter transit-amplifying divisions (b). Neurogenic astrocytes maintain normal astrocyte morphology even as they have upregulated *Ascl1* and initiated DNA synthesis in preparation for transit-amplifying divisions (c, 1-cell stage. Note that EdU, which was administered for the last two weeks prior to sacrifice, has been incorporated into the DNA of this single astrocyte. This indicates that astrocytes initiate S phase while still retaining astrocyte features.). As transit-amplifying divisions occur, astrocytic processes are gradually lost by the dividing cells (c, “2-10 cells”. See also Movie S1.). The astrocyte marker S100β is not expressed by neurogenic-lineage cells in the SVZ, neither in single Ascl1$^+$ cells or clustered transit-amplifying cells (d, e [red arrowheads]; each data point in (d) is one cell). In contrast, many striatal astrocytes express high levels of S100β (d, f [blue arrowheads]). Consequently, many (but not all) striatal transit-amplifying cells retain low amounts of residual S100β protein (d, g [blue arrowheads]) as remnants of their origin as parenchymal astrocytes. Thus, lingering S100β protein can be used as a short-term lineage-tracing marker for transit-amplifying cells derived from striatal astrocytes. (LV, lateral ventricle). Scale bar (c): 10 µm

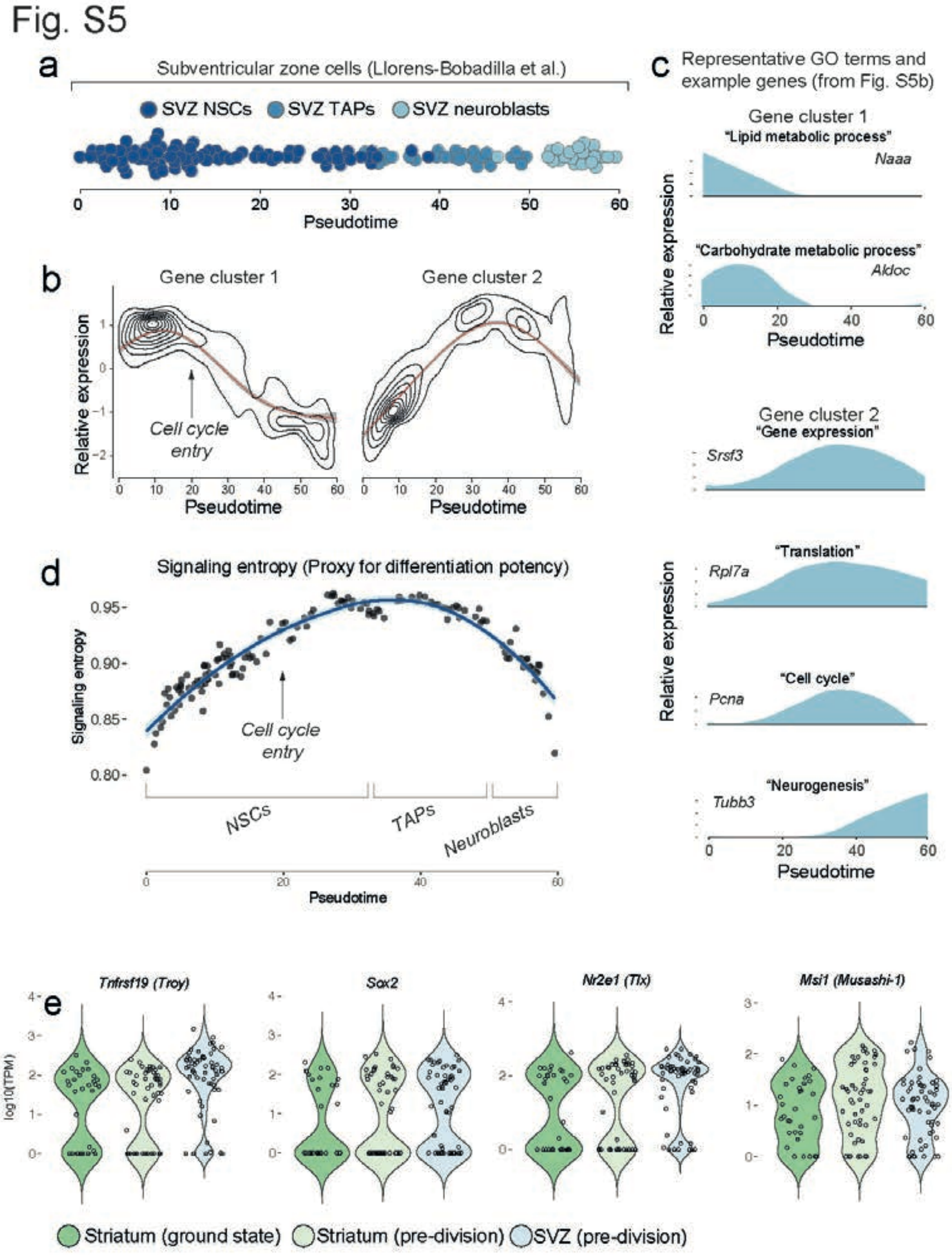


**Figure S5. Gene clustering analysis of SVZ neural stem cells and their progeny from the dataset published by Llorens-Bobadilla et al.** We analyzed a previously published dataset of SVZ neural stem cells and their progeny. Monocle reconstructs SVZ neurogenesis along pseudotime (a; cells are colored based on cell groups defined in Llorens-Bobadilla et al.). An analysis using Monocle's gene clustering algorithm shows that, as SVZ stem cells enter transit-amplifying divisions, genes associated with lipid and carbohydrate metabolism are downregulated whereas genes associated with gene expression are upregulated (b-c). Signaling entropy, an approximation of a cell's differentiation potency, increases as SVZ stem cells are activated and initiate transit-amplifying divisions (d). Some classical markers of neural stem cells are already expressed by ground-state striatal astrocytes and their RNA levels do not change after *Rbpj* deletion (e).

### Fig. S6

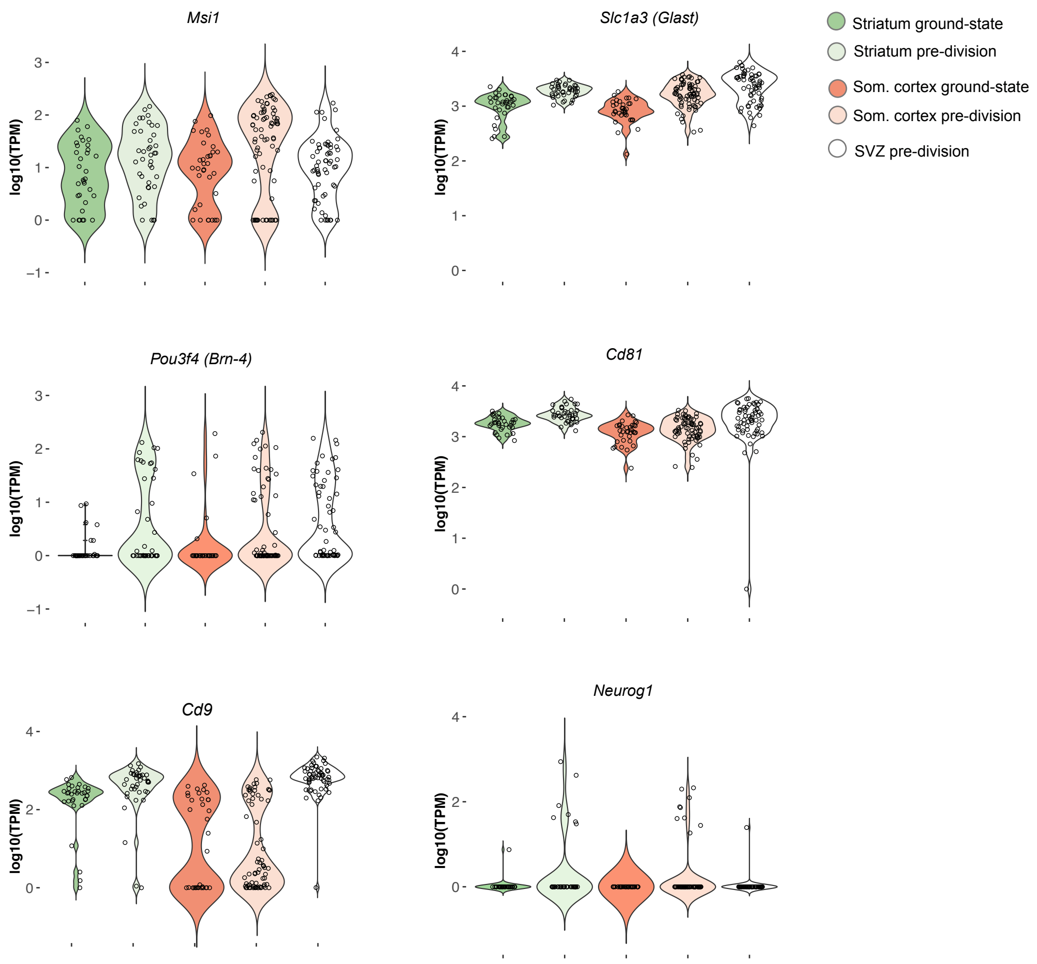


**Figure S6. Expression levels of neural stem cell markers in striatum, somatosensory cortex and SVZ.** In the pre-division stage, cortical astrocytes have upregulated many neurogenesis-associated genes to the same levels as in the striatum and SVZ. One neural stem cell marker, *Cd9*, was conspicuously lower in the cortical astrocytes than in the striatum and SVZ. Another gene, *Neurog1*, was uniquely upregulated by the parenchymal astrocytes and was not expressed at any differentiation stage in the SVZ.

Fig. S7

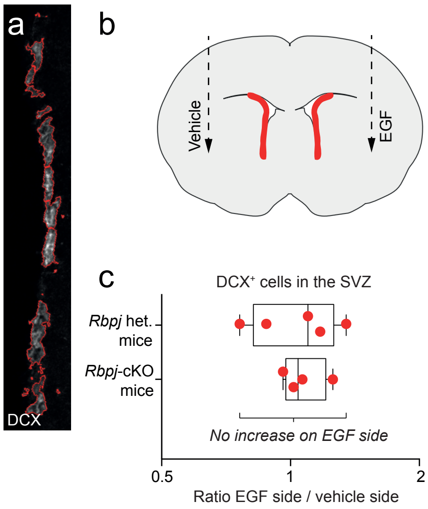


**Figure S7. EGF injection into the lateral striatum does not change the amount of SVZ neuroblasts**. Dcx signal intensity was used as a proxy for Dcx cell numbers in the SVZ. Dcx signal intensity was quantified using CellProfiler, which delineates $Dcx^+$ cells and measures their signal intensity (red outlines in a). In mice injected with EGF into one striatum and a vehicle solution in the other striatum (b), we could not detect any change in the amount of SVZ neuroblasts in the EGF-injected side, neither in mice heterozygous or homozygous for the *Rbpj* mutation. This is shown in (c); here, each red dot represents one animal, and the position of each dot indicates whether that mouse had more SVZ neuroblasts on the vehicle side or the EGF side. On average, there is no significant difference between the vehicle side and the EGF side, neither in *Rbpj*-heterozygous or *Rbpj*–KO mice. This indicates that EGF injection in the lateral striatum does not significantly affect the number of SVZ neuroblasts.

**Table S1. Genes differentially expressed between ground-state and *Rbpj*-deficient (two weeks post-tamoxifen) astrocytes.** Data is shown for both striatum and somatosensory cortex.

**Table S2. Lists of genes characteristic for the four gene clusters in Fig. 2e.**

**Table S3. Details about mice transplanted with neural stem cells (Fig. 6l-n).**
